# Requirement for STAT3 and its target, TFCP2L1, in self-renewal of naïve pluripotent stem cells *in vivo* and *in vitro*

**DOI:** 10.1101/2022.09.23.509173

**Authors:** Sophie Kraunsoe, Takuya Azami, Yihan Pei, Graziano Martello, Kenneth Jones, Thorsten Boroviak, Jennifer Nichols

**Affiliations:** Wellcome Trust – Medical Research Council Stem Cell Institute, University of Cambridge, Jeffrey Cheah Biomedical Centre, Puddicombe Way, Cambridge CB2 0AW, UK; Department of Physiology, Development and Neuroscience, University of Cambridge, Tennis Court Road, Cambridge CB2 3EG, UK; Centre for Trophoblast Research, University of Cambridge, UK; MRC Human Genetics Unit, Institute of Genetics and Cancer, University of Edinburgh, Western General Hospital, Crewe Road South, Edinburgh EH4 2XU, UK; School of Bioscience, University of Nottingham Sutton Bonington Campus, Loughborough, LE12 5RD, UK; Department of Biology, University of Padua, Italy; Department of Biochemistry, University of Cambridge

**Keywords:** Naïve pluripotency, Embryonic diapause, STAT3, TFCP2L1, Blastocyst, Embryonic stem cells

## Abstract

We previously demonstrated gradual loss of epiblast during diapause in embryos lacking components of the LIF/IL6 receptor. Here we explore requirement for the downstream signalling transducer and activator of transcription, STAT3 and its target, TFCP2L1, in maintenance of naïve pluripotency. Unlike conventional markers, such as NANOG, which remains high in epiblast until implantation, both STAT3 and TFCP2L1 proteins decline during blastocyst expansion, but intensify in the embryonic region after induction of diapause, as observed visually and confirmed using our novel image analysis tool, consistent with our previous transcriptional expression data. Embryos lacking STAT3 or TFCP2L1, underwent catastrophic loss of most of the inner cell mass during the first few days of diapause, implicating involvement of signals in addition to LIF/IL6 for sustaining naïve pluripotency *in vivo*. By blocking MEK/ERK signalling from the morula stage we could derive embryonic stem cells with high efficiency from STAT3 null embryos, but not those lacking TFCP2L1, suggesting a hitherto unknown additional role for this essential STAT3 target in transition from embryo to embryonic stem cells *in vitro*.

**Summary Statement:** Inducing diapause in mouse embryos demonstrates that STAT3 and TFCP2L1 are essential for self-renewal of the epiblast, but only TFCP2L1 is required for derivation of embryonic stem cells.

## Introduction

Embryonic stem cell lines (ESCs) derived from epiblasts of preimplantation mouse embryos (Evans and Kaufman, 1981; Martin, 1981) have been used extensively to study and model mammalian development, since they can be expanded in culture, whilst retaining the ability to differentiate into all tissues of the body. This flexible state is known as ‘naïve pluripotency’ (Nichols and Smith, 2009). ESCs can self-renew in medium supplemented with leukaemia inhibitory factor (LIF) (Smith et al., 1988; Williams et al., 1988), operating via signal transducer and activator of transcription (STAT)3 (Burdon et al., 1999; Matsuda et al., 1999; Niwa et al., 1998). The LIF receptor complex comprising LIFR and gp130 (also known as IL6ST) activates Janus-associated kinases (JAKs), which phosphorylate STAT3 at tyrosine 705 (pY705) (Ni et al., 2004; Zhang et al., 2000). Mutation of pY705 ablates ESC self-renewal in standard (serum/LIF) culture conditions (Huang et al., 2014); LIF or related cytokines were therefore previously considered essential for ESC self-renewal. However, a refined serum-free culture regime, ‘2i’, based upon dual inhibition of Glycogen Synthase Kinase (GSK)3 and MEK/ERK, allows ESC derivation from STAT3 mutant embryos by incubation from the morula stage (Ying et al., 2008). STAT3 null ESCs enabled analysis of downstream signalling events associated with ESC self-renewal, and thus characterisation of the signalling network operative during maintenance of naïve pluripotency *in vitro* (Martello et al., 2013; Ye et al., 2013). The most significant player emerging from this analysis was TFCP2L1 (also known as CRTR-1) whose forced expression could rescue STAT3 null ESCs in serum/LIF culture. Using pathway analysis and computational modelling an essential role for TFCP2L1 in ESC maintenance was proposed and supported experimentally (Dunn et al., 2014). Moreover, transfection of *Tfcp2l1* into epiblast stem cells (EpiSCs) derived from postimplantation epiblasts (Brons et al., 2007; Tesar et al., 2007) could direct reprogramming from primed to naïve pluripotency, confirming participation of TFCP2L1 in the naïve pluripotency network (Martello et al., 2013; Ye et al., 2013).

Combined maternal and zygotic deletion revealed an essential requirement for STAT3 during blastocyst expansion, confirming its suspected function in epiblast formation (Do et al., 2013). However, perdurance of maternal STAT3 protein in zygotic null embryos permits developmental progression beyond cleavage stages, allowing them to implant in the uterus, where they acquire abnormalities (Takeda et al., 1997). Interestingly, mutation of either LIFR or gp130 results in considerably less severe phenotypes (Li et al., 1995; Yoshida et al., 1996), supporting previous suggestions of additional alternative requirement for STAT3 signalling in early mouse development (Kristensen et al., 2005).

In normal laboratory rodents, the state of naïve pluripotency is relatively transient, lasting no more than two days. It is therefore debatable whether self-renewal of naïve pluripotent stem cells occurs at this stage during uninterrupted development. Conveniently, murine preimplantation embryogenesis can be prolonged by diapause, a natural, facultative phenomenon ensuing when a dam conceives whilst suckling a previous litter. Embryos progress to the periimplantation stage, embryonic day (E)4.5, but delay implantation until a source of oestrogen is regained. Diapause can be achieved experimentally by ovariectomy prior to the physiological burst of oestrogen secretion at E2.5 (Weitlauf and Greenwald, 1968). Healthy blastocysts are able to sustain diapause for more than a month, then resume normal development (Arena et al., 2021). In previous studies, we showed that epiblast in diapause embryos lacking LIFR or gp130 was gradually reduced by apoptosis, leaving only trophectoderm and primitive endoderm (PrE) (Nichols et al., 2001). Loss of STAT3 or its target, TFCP2L1, may be anticipated to result in a more dramatic phenotype. Here we show that both STAT3 and TFCP2L1 are critically required for diapause, with almost complete loss of both epiblast and PrE comprising the inner cell mass (ICM) occurring within 4 days of diapause onset (6 days after ovariectomy). Although both pY705 STAT3 and TFCP2L1 are downregulated in the epiblast before implantation, their levels intensify during diapause, consistent with previously published RNA profiling (Boroviak et al., 2015). To enhance optical compartmentalisation of tissue types in periimplantation and diapause embryos, we developed an image analysis tool to quantify confocal readout of protein levels and distribution for pY705 STAT3 and TFCP2L1. Despite the rapid loss of epiblast in diapaused mutant embryos, we show that capture of self-renewing ESCs from STAT3 null embryos in culture is as efficient as from wild type and heterozygous embryos. In contrast, no ESCs could be derived from embryos lacking TFCP2L1, implicating alternative functions for this factor in replication of naïve pluripotent epiblast cells that is not directed by STAT3 signalling.

## Results

### STAT3 pY705 and TFCP2L1 peak in epiblast during embryonic diapause

The potential roles of STAT3 and TFCP2L1 in sustaining pluripotency *in vivo* were investigated via induction of diapause (Fig.1A), as described previously (Nichols et al., 2001). Bilateral ovariectomy was performed on female CD1 mice 2.5 days after mating by males of the same strain. Embryos were flushed 6 days later, and were therefore in the state of diapause for 4 days. Immunofluorescence (IF) using antibodies raised against NANOG, STAT3 (pY705) and TFCP2L1 was performed using our standard protocol (Silva et al., 2009), but with methanol permeabilization, on non-diapaused embryos at E3.5 and 4.5 and following 4 days of diapause (Fig.1B). Quantification for levels of NANOG, pY705 and TFCP2L1 proteins in the embryonic region was enabled by manual cropping to exclude the abembryonic, mural trophectoderm region (Fig.2A), and application of a novel image analysis tool (Fig.2B). In concordance with visual appearance of the confocal images, a significant increase of both pY705 and TFCP2L1 IF was quantified in diapause epiblast compared with non-diapaused periimplantation embryos at E4.5 (Fig.2C), consistent with mRNA levels and the transcriptional resemblance of diapause epiblast to self-renewing ESCs *in vitro* (Boroviak et al., 2015). During manual cropping of the images analysed with the image analysis tool, most mural trophectoderm was removed to prevent these cells confounding the analysis. By creating spatially reconstructed *in silico* embryos, it can be observed that cells selected for high levels of STAT3 pY705 or TFCP2L1 are enriched in the epiblast compartment of diapaused embryos (Fig.3A,B). In contrast, E4.5 embryos have lower levels of STAT3 pY705 in their epiblast compartment compared to E3.5 and diapaused embryos (Fig3C).

**Figure 1.**
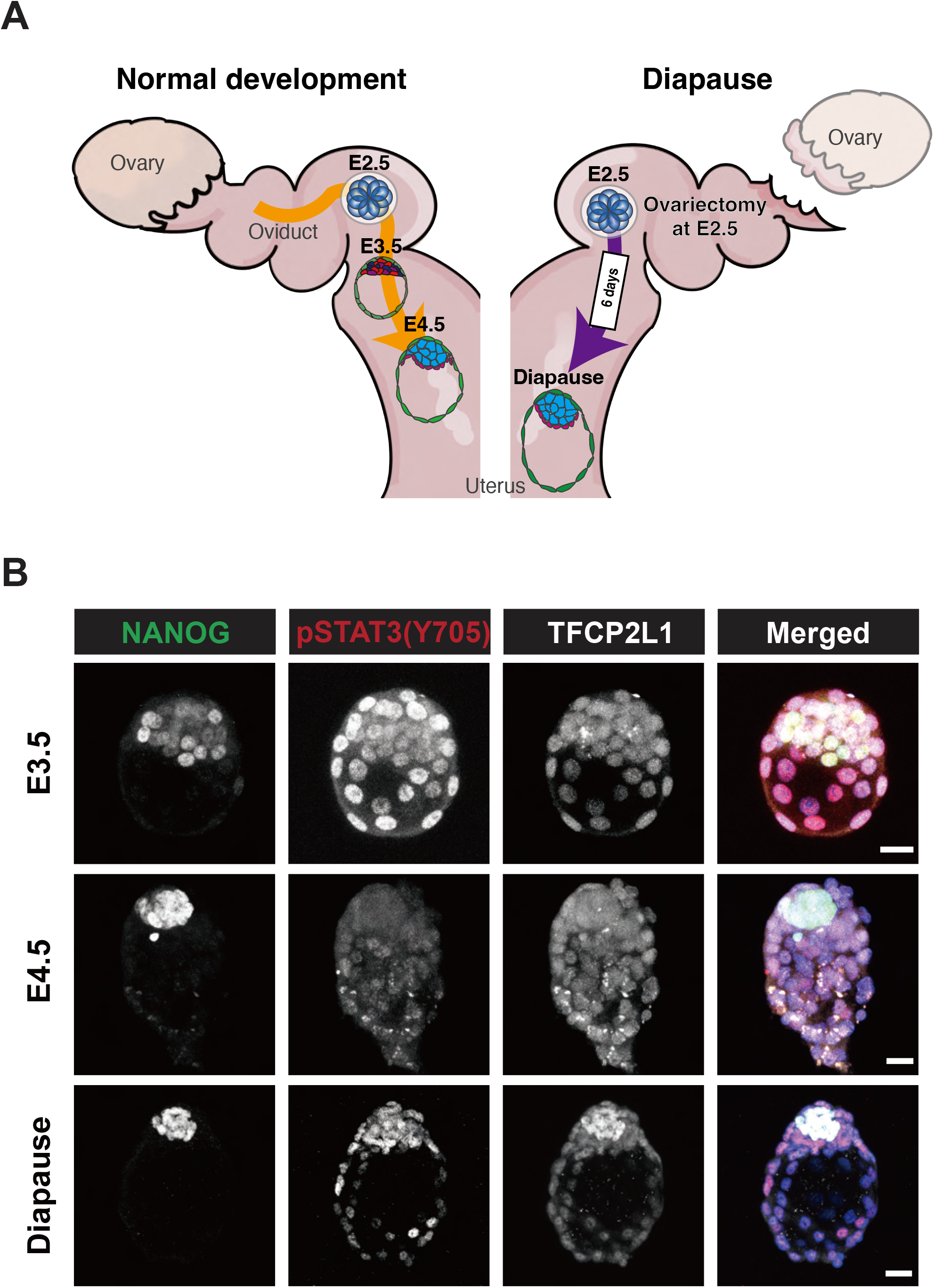
Distribution of pSTAT3(Y705) and TFCP2L1 in preimplantation and diapause embryos. (A) Schematic of mouse reproductive system representing normal preimplantation development (left) and induction of diapause (right). Diapause was induced by surgical removal of both ovaries at E2.5 (before the physiological burst of oestrogen production) and embryos were collected 6 days later, thus being diapaused for 4 days. (B) Confocal images of NANOG, pSTAT3(Y705) and TFCP2L1 IF in preimplantation E3.5, 4.5 and diapause embryos. Scale bar = 20 µm.

**Figure 2.**
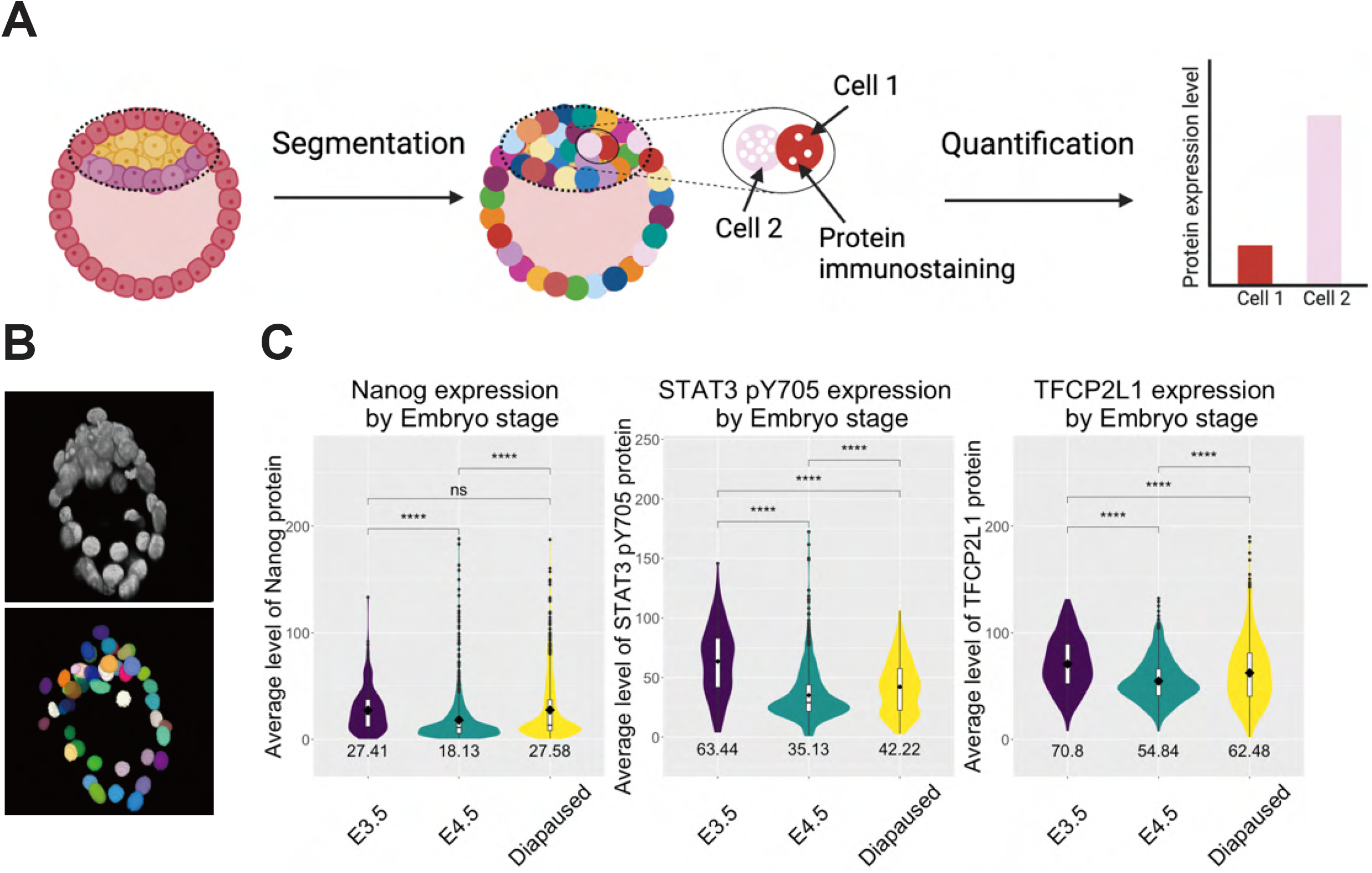
Development of analysis tool for pre-implantation mouse embryo IF. (A) Schematic of image analysis pipeline to segment individual nuclei in 3D and quantify fluorescence of each channel. The region of each embryo to be quantified is depicted by a dotted red oval, thus excluding the mural trophectoderm region that does not participate in subsequent formation of the foetus. (B) Representative example embryos from each developmental stage with nuclei represented by a scatter point, and colour coded based on (STAT3)pY705 or TFCP2L1 integrated density. (C) Violin plots with overlaid boxplots of the integrated density of pY705 and TFCP2L1 expression across each nucleus from each embryo by stage (E3.5, n = 29, E4.5, n= 33, diapause, n = 54). Segmented nuclei were filtered for DAPI signal and volume to remove any erroneous segmentation. * *P* < 0.01.

**Figure 3.**
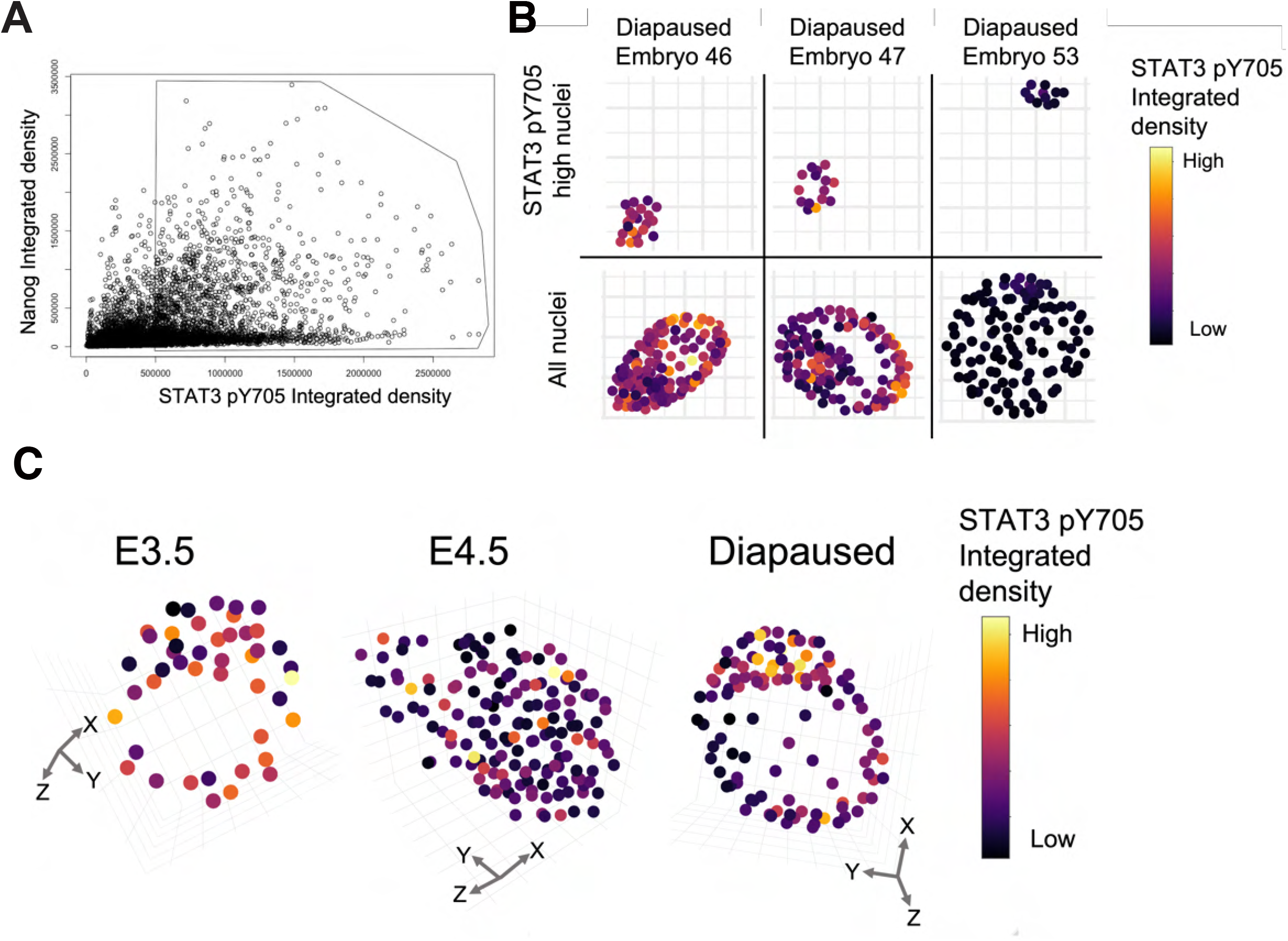
Example of analysis of embryo IF. (A) Nuclei are colour-coded by stage dispersed in a 3D co-expression space based on integrated density (sum across all pixel values in the 3D object) of NANOG and pY705 and in each segmented nucleus. The polygon selection encloses cells with a pY705 raw integrated density above 5 × 10^5^. (B) Distribution of the selected high pY705 cells within the polygon from (A) in their embryo of origin for 3 representative diapaused embryos (top) with all nuclei shown below. (C) Representative example embryos from each developmental stage with nuclei represented by a scatter point and colour coded based on pY705 integrated density.

### Propagation of naïve pluripotency in vivo requires STAT3 and TFCP2L1

To assess functionality of STAT3 signalling during maintenance of naïve pluripotency, heterozygous mice were mated *inter se* to generate wild type (WT), heterozygous (het) and mutant (null) embryos for *Stat3* or *Tfcp2l1*. Diapause embryos were recovered 6 days after ovariectomy and IF for NANOG, pY705 and TFCP2L1 was performed. Whereas WT and het embryos possessed large ICMs with many NANOG, pY705 and TFCP2L1 positive cells, embryos lacking STAT3 or TFCP2L1 exhibited complete loss or severe reduction of the whole ICM (Fig.4A,B). In both cases, null embryos were underrepresented (Fig.4C), probably attributed to loss before retrieval owing to catastrophic reduction of the ICM impacting on trophectoderm expansion and embryo integrity. Taken together, these results implicate immediate requirement for STAT3 signalling and functional TFCP2L1 during diapause.

**Figure 4.**
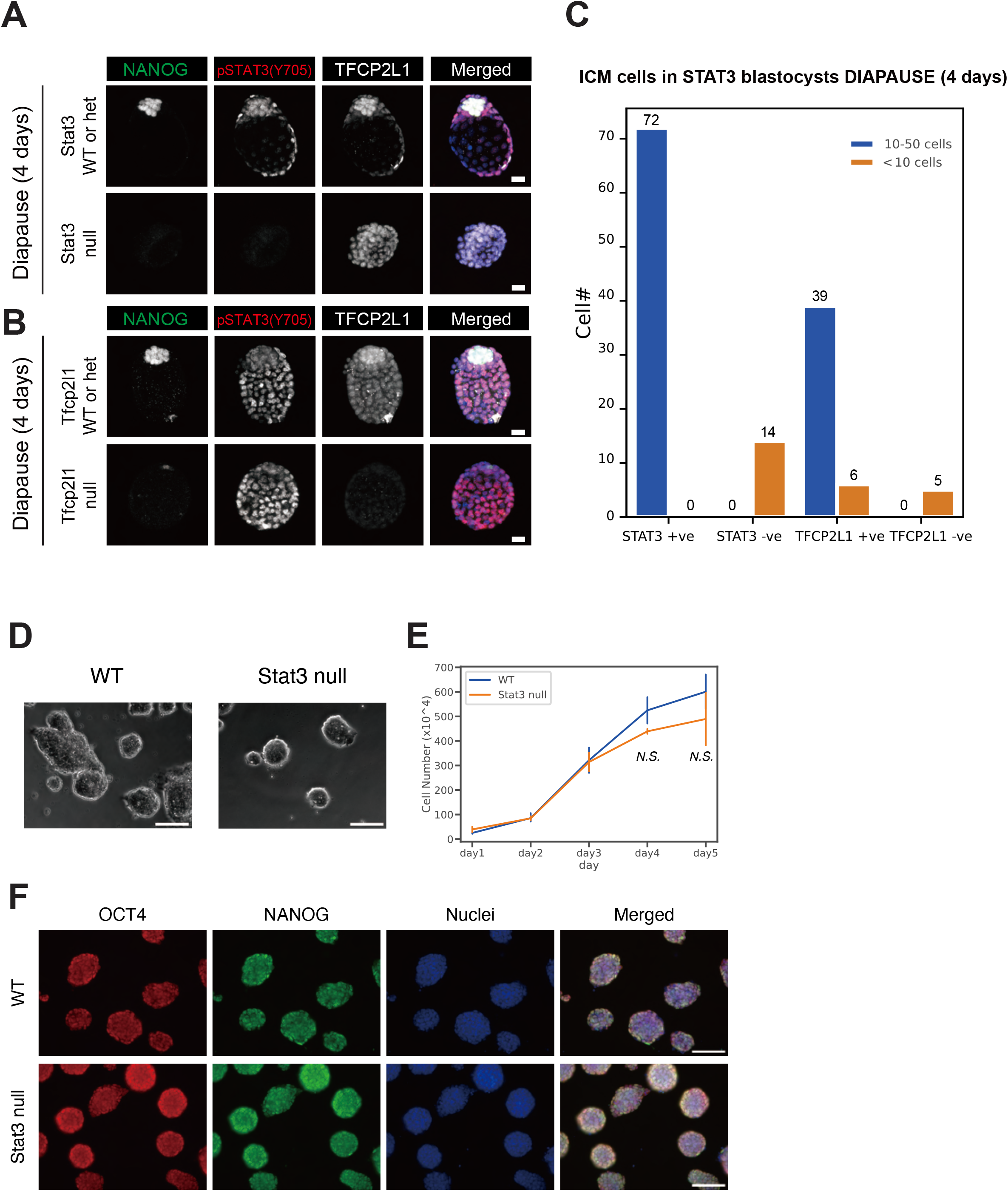
STAT3 and TFCP2L1 are required for ICM maintenance during diapause, but STAT3 is not required for ESC derivation or self-renewal. (A, B) IF of NANOG, pY705 and TFCP2L1 in diapause for (A) *Stat3* WT/het and null embryos and (B) *Tfcp2l1* WT/het and null embryos. Scale bar = 20 µm. (C) Number of ICM cells in *Stat3* or *Tfcp2l1* WT/Het and null diapause embryos. (D) Bright field images of WT and *Stat3* null ESCs. Scale bar = 100 µm. (E) Proliferation based on cell number counts of WT and *Stat3* null ESCs cultured in N2B27+2i medium for 5 days (day4: p=0.14, day5: p=0.55 by Student’s T test). (F) Immunofluorescence of OCT4 and NANOG in WT and *Stat3* null ESCs. Scale bar = 100 µm.

### Capture of ESCs from embryos

Our previous derivation protocol based upon blocking MEK/ERK and GSK3 from the morula stage of development (Ying et al., 2008) was used to capture CD1 background ESCs. 41 WT or het and 13 null ESC lines were generated from 56 embryos by *inter se* mating of *Stat3* het mice (Table 1). Each embryo was genotyped by PCR using lysed trophectoderm produced during immunosurgery (Nichols et al., 1998; Solter and Knowles, 1975). *Stat3* null lines were confirmed by genotyping after expansion. *Stat3* null and WT cell lines could be propagated indistinguishably in 2i medium (Fig.4D) and no distinct difference in cell cycle kinetics could be perceived between them (Fig.4E). IF for OCT4 and NANOG confirmed naïve pluripotent identity for both WT and STAT3 null ESC lines (Fig.4F). Conversely, from 65 morulae generated from *Tfcp2l1* het inter-cross, 6 WT and 49 het, but no null ESC lines were derived from 65 embryos (Table 1), suggesting a distinct requirement for TFCP2L1 in capture of pluripotency *in vitro*.

**Table 1.**
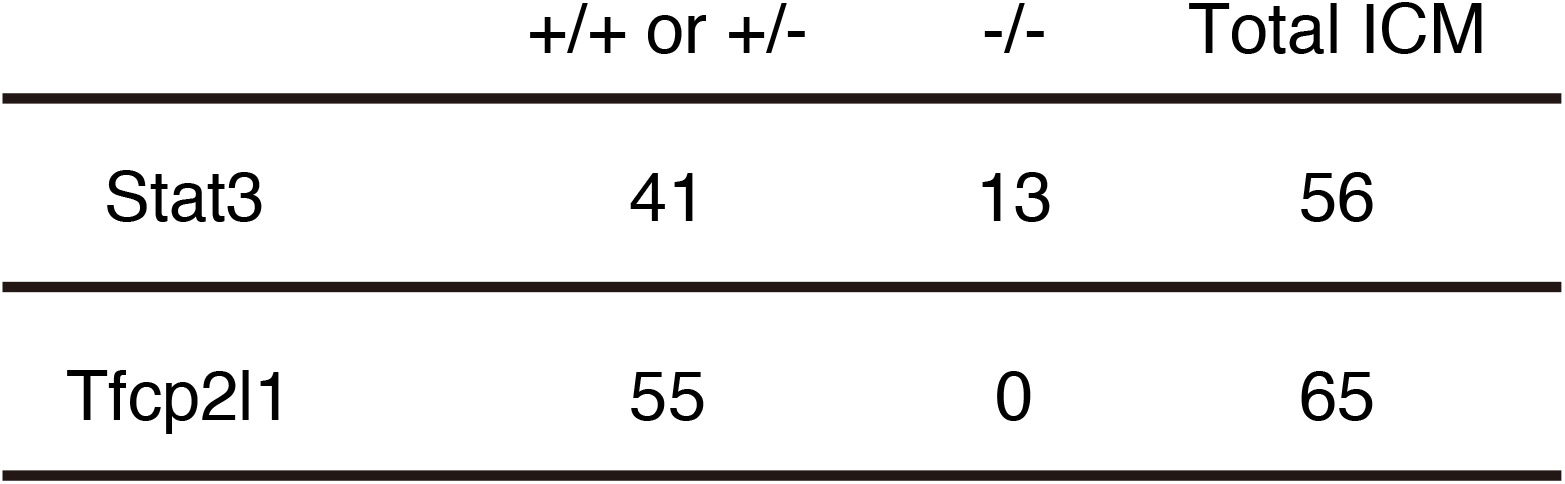
ESCs derivation from ICM of Stat3 or Tfcp2l1 heterozygous intercrosses

## Discussion

Derivation of *Stat3* null ESCs previously facilitated interrogation of STAT3 targets and highlighted TFCP2L1 as the most significant *in vitro* (Martello et al., 2013; Ying et al., 2008). To investigate the potential role of STAT3 and TFCP2L1 in maintenance of naïve pluripotency *in vivo* we induced embryonic diapause. IF for pSTAT3(Y705), the protein product utilised for self-renewal of ESCs (Huang et al., 2014), and TFCP2L1 both become visibly enriched in the epiblast during diapause (Fig.1). This observation was validated using a novel tool developed for quantification of IF images of tightly compacted nuclei (Figs.2-3), thus implying a possible role for these factors for epiblast self-renewal *in vivo*. During diapause, in contrast to the phenotype observed following deletion of LIFR or its co-receptor, gp130, which resulted in gradual loss of epiblast, but not PrE (Nichols et al., 2001), embryos lacking either STAT3 or TFCP2L1 lost virtually the entire ICM within only a few days (Fig.4). This more dramatic phenotype may be a consequence of the precipitous depletion of epiblast, the source of PrE-inducing FGF4 (Yamanaka et al., 2010), compared with deletion of LIF receptor complex components (Nichols et al., 2001). STAT3 also promotes anti-apoptotic activity (Hirano et al., 2000), which could contribute to the enhanced PrE population reported in blastocysts supplemented with IL6 (Anderson et al., 2017; Morgani and Brickman, 2015). We conclude that STAT3 signalling is essential to maintain naïve pluripotency *in vivo* and operates as a signal transducer for pathways in addition to that induced by IL6 family cytokines.

Interestingly, we recently found precocious expression of PrE markers such as *Pdgrfa, Sox17* and *Gata4* in *Stat3* null embryos at the mid blastocyst stage (E3.5), whereas emerging epiblast cells at E3.75 prematurely activated the postimplantation epiblast genes *Utf1, Otx2* and its targets *Dnmt3a and b* (Betto et al., 2021), which presumably instigated reduction of FGF4 secretion. However, previous observations of PrE persistence during diapause when LIFR or gp130 are deleted (Nichols et al., 2001) imply independence of this branch of STAT3 signalling for PrE maintenance. The present data implicate STAT3 signalling, via TFCP2L1, in PrE maintenance *in vivo*. Our failure to derive ESCs from *Tfcp2l1* null embryos using the strategy that proved highly successful for generation of *Stat3* null ESCs (Table 1) implies an unexpected STAT3-independent role for TFCP2L1 in transition towards *in vitro* self-renewal of pluripotent stem cells. TFCP2L1 plays a role in upregulation of *Nanog* (Ye et al., 2013), which is also indispensable for derivation of ESCs from mouse embryos (Silva et al., 2009). Interestingly, both *Nanog* and *Tfcp2l1* can be deleted from established ESCs cultured in 2i/LIF (Chambers et al., 2007; Martello et al., 2013; Yan et al., 2021), implicating compensation by the robust and redundant network of pluripotency factors assembled in ESCs *in vitro* (Dunn et al., 2014). This pathway connection may explain the failure of embryos lacking TFCP2L1 to yield ESCs and further highlights the distinct requirements for self-renewal of naïve pluripotent cells *in vivo* compared with established cell lines *in vitro*.

## Materials and Methods

### Mice, husbandry and embryos

Experiments were performed in accordance with EU guidelines for the care and use of laboratory animals and under the authority of appropriate UK governmental legislation. Use of animals in this project was approved by the Animal Welfare and Ethical Review Body for the University of Cambridge and relevant Home Office licences are in place.

Mice were maintained on a lighting regime of 12:12 hours light:dark with food and water supplied *ad libitum. Stat3* mice heterozygous for replacement of exons 20-22 with Neomycin resistance (Takeda et al., 1997) were backcrossed to CD1 mice. *Tfcp2l1* heterozygous mice were generated from ESCs targeted using CRISPR strategy in E14 ESCs obtained from Jackson Labs Knockout Mouse Project (KOMP), via injection into C57BL/6 blastocysts to generate chimaeras. Male chimaeras were mated with CD1 females; grey pups were genotyped by PCR of ear biopsies and robust males selected for further backcrossing to CD1 females. Both STAT3 and TFCP2L1 mouse lines were maintained by backcrossing to CD1. Embryos were generated from *Stat3*^+/-^ or *Tfcp2l1*^*+/-*^ *inter se* natural mating. Detection of a copulation plug in the morning after mating indicated embryonic day (E)0.5. Embryos were isolated in M2 medium (Sigma-Aldrich).

### Genotyping

Mice were genotyped by PCR using ear biopsies collected within 4 weeks of birth and genomic DNA was extracted using Extract-N-Amp tissue prep kit (Sigma-Aldrich). Embryos were genotyped using either immune-reactivity to antibody raised against either STAT3 pY705 or TFCP2L1 in the case of those imaged for confocal analysis, or PCR analysis of trophectoderm lysate for ESC derivation. Amplification was carried out on 5 µL of lysate for 35 cycles (following 95°C hot start for 10 minutes) of 94°C, 15 seconds; 60°C, 12 seconds; 72°C, 60 seconds, with a final extension at 72°C for 10 minutes. Reaction products were resolved by agarose gel electrophoresis. Primers used for genotyping PCR are listed in Table.S1.

### Induction of diapause

To determine the requirement for STAT3 or TFCP2L1 during maintenance of the embryo during delayed implantation, CD1 females or het females mated by het males were surgically ovariectomised under general anaesthesia as described previously (Nichols et al., 2001) before the embryos reached E2.5. Diapause embryos were flushed 6 days later and fixed for IF.

### Derivation and culture of ESC lines

Morulae were collected from het females 2.5 days after mating by het males and used for ESC derivation as described previously (Ying et al., 2008) by culture to the blastocyst stage in KSOM supplemented with 2i, consisting of 1 µM PD0325901 and 3 µM CHIR99021, transfer of ICMs isolated by immunosurgery (Solter and Knowles, 1975) to 48-well plates containing 2i in N2B27 medium, one per well. WT and *Stat3* null ESCs were expanded and maintained in N2B27 supplemented with 2i or 2i/LIF on gelatin-coated plates at 37°C in 7% CO_2_ and passaged by enzymatic disaggregation every 2-3 days.

### Immunofluorescence (IF)

Embryos were fixed in 4% paraformaldehyde (PFA) for 30 min at room temperature (RT), followed by washing in 0.5% polyvinylpyrrollidone (PVP) in PBS. Embryos were permeabilized in 0.5% Triton X-100 in PBS for 15 min and blocked with 2% donkey serum, 2.5% BSA, and 0.1% Tween20 in PBS (blocking solution) for 1 h at RT. For phosphorylated-STAT3 staining, permeabilization was performed in absolute methanol for 10 min at −20°C. Primary antibodies were diluted in blocking solution and embryos incubated in these overnight at 4°C. After washing in 0.1% Tween 20 in PBS, embryos were incubated with Alexa Fluor-conjugated secondary antibodies (Thermo) at 1:500 dilution in blocking solution for 1 h at RT. Nuclear staining was carried out with Hoechst33342 or DAPI (Thermo). Primary and secondary antibodies used are listed in Table S2.

### Imaging and image analysis of embryos

Images for embryos were acquired using TCS SP5 (Leica) confocal microscope and processed with ImageJ. Quantification of IF images was achieved using a computational pipeline to extract data on the intensity of antibody staining in individual nuclei. The 2D StarDist segmentation Fiji plugin (parameters: ‘percentileBottom’:‘1.0’, ‘percentileTop’:‘99.9’, ‘probThresh’:‘0.2’, ‘nmsThresh’:‘0.2’) was used to segment the DNA channel of IF images before using the Trackmate plugin to ‘track’ objects through the z-stack to generate 3D objects (parameters: area threshold > 5, <30, track duration > 5) (Ershov et al., 2022). An unsharp mask was applied to the DNA channel prior to segmentation (parameters: radius = 15, mask = 0.6). Nuclear segmentation of embryos was used as a mask to measure fluorescence intensity, xyz position, and morphological parameters (e.g., volume) of each nucleus (Ollion et al., 2013; Pietzsch et al., 2015). Assessment of the histograms for signal intensity in the DNA channel and nuclear volume allowed thresholds to be set to remove erroneously segmented nuclei. Violin plots were generated using the ggplot2 package in R and statistical significance between embryo groups was assessed via pair-wise unpaired student’s t-tests. Nuclei were visualised in a 2D gene expression space based on NANOG and STAT3 intensity and selected nuclei were plotted back into the ‘in silico’ embryos, based on their positional information and colour coded by STAT3 expression levels. ‘In silico’ embryos were generated using xyz centroid information from the segmentation mask and colour coded according to STAT3 intensity.

## Acknowledgements

We wish to thank William Mansfield for generation of Tfcp2l1+/-mouse line; Shizuo Akira for providing Stat3+/-mice; Peter Humphreys and Darran Clement for advanced imaging and analysis; UBS, Cambridge for animal husbandry.

No competing interests declared

## Data Availability

Fiji macro and R codes will be made available at https://github.com/SKraunsoe/STAT3_embryo_segmentation_analysis

